# Co-infection of *Candidatus* Piscichlamydia trichopodus (order Chlamydiales) and *Henneguya* sp (Myxosporea, Myxobolidae) in snakeskin gourami *Trichopodus pectoralis* (Regan 1910)

**DOI:** 10.1101/2021.06.01.446678

**Authors:** Nguyen Dinh-Hung, Ha Thanh Dong, Chayanit Soontara, Channarong Rodkhum, Sukkrit Nimitkul, Prapansak Srisapoome, Satid Chatchaiphan, Pattanapon Kayansamruaj

## Abstract

The present study describes a simultaneous infection of a novel *Chlamydia*-like organism (CLO) with a Myxozoa parasite, *Henneguya* sp. in snakeskin gourami *Trichopodus pectoralis* in Thailand. A new CLO is proposed “*Candidatus* Piscichlamydia trichopodus” (CPT) based on 16S rRNA phylogenetic analysis. Systemic intracellular CPT infection was confirmed by histological examination, *in situ* hybridization, PCR assay, and sequencing of 16S rRNA. This novel pathogen belongs to the order *Chlamydiales* but differs in certain aspects from other species. The histopathological changes associated with CPT infection were different from the typical pathological lesions of epitheliocystis caused by previously known CLO. Unlike other CLO, CPT localized in the connective tissue rather than in the epithelial cells and formed smaller clumps of intracellular bacteria that stained dark blue with hematoxylin. On the other hand, typical myxospores of the genus *Henneguya* with tails were observed in the gill sections. Infection with *Henneguya* sp. resulted in extensive destruction of the gill filaments, most likely leading to respiratory distress. Due to the frequency of co-infections and the unavailability of culture methods for CLO and *Henneguya* sp, it was difficult to determine which pathogens were directly responsible for the associated mortality. However, co-infections may increase the negative impact on the host and the severity of the disease. Given the commercial importance of the snakeskin gourami and its significant aquaculture potential, the findings of this study are important for further studies on disease prevention.

## 1. Introduction

Snakeskin gourami, *Trichopodus pectoralis*, is native to Southeast Asia and commonly found in the Mekong and Chao Phraya basins of Cambodia, Thailand, Southern Vietnam, and Laos [1]. In Thailand, the snakeskin gourami is a highly economic species and has become one of the five most important freshwater species in aquaculture [2, 3]. Increased incidence of parasitic and bacterial diseases is one of the major obstacles to the farming of this species [4]. However, studies on the occurrence of diseases in *T. pectoralis* are still very scarce [5]. The pathogen fauna of this species is poorly understood and may contain pathogens that have not yet been described in the literature.

Myxosporeans are diverse and widespread parasites that cause severe economic damage to fish worldwide [6-8]. The genus *Henneguya* includes more than 200 species and is one of the most diverse myxosporean genera in the family *Myxobolidae* [8]. Some species of the genus *Henneguya* are responsible for diseases leading to high mortality rates, but most species are thought to have little or no negative impact on fish health [6, 9]. Infection from *Henneguya* sp. usually occurs in the gills and is characterized by the presence of cyst-like structures on the gill filaments [6, 7, 10]. Infection can devastate fish populations when the parasites multiply in high densities on the gills and leads to respiratory failure, especially in juvenile fish [11, 12]. Other commonly known pathogens that cause gill cysts in fish are bacteria from the order *Chlamydiales*. These bacteria typically cause epithelial cysts in the skin and gills called epitheliocystis [13-15]. However, it is noteworthy that different *Chlamydia*-like organisms (CLOs) have been found to have different pathology upon infection, possibly depending on the chlamydial species, the infected host, and the affected tissue [16]. To date, unavailability of culture techniques for chlamydial pathogens has been a major obstacle for *in vitro* studies [13, 16].

In the present study, we described for the first time a systemic pathology caused by a novel *Chlamydia*-like organism, *Candidatus* Piscichlamydia trichopodus (order *Chlamydiales*) and a gill parasite *Henneguya* sp. (Myxosporea, Myxobolidae) infecting the same fish population. Pathogen characterization was performedbased on molecular analyses with detailed histopathological observations and confirmation by *in situ* hybridization (ISH).

## 2. Materials and methods

### 2.1. Fish and case history

In March 2021, higher than average mortality was observed in snakeskin gourami fingerlings in two nursery ponds in central Thailand. History entails that snakeskin gourami fingerlings (0.3g ± 0.05) reared by traditional methods [17] were purchased from a hatchery. Male and female broodstock were naturally mated in an earthen pond. Eggs were spawned in natural bubble nests made by male gourami in the same pond. Offspring were harvested at the size of 3.0 cm (weight 0.2-0.3 g) and 150,000 fish were delivered to the two ponds mentioned above. The fish were kept in a 5 m^2^ net in the pond and fed daily with a commercial pellet diet containing 28% protein (Betagro, Thailand), administered at a rate of 3% of body weight. Mortality was recorded on the second day after the fry were introduced. Daily mortality recorded from 1 to 5 days after onset of disease (dao) were 20, 50, 150, 200, and 400 fish, respectively. Affected fish showed lethargy, gasping at the water surface and loss of appetite followed by mortality. At 3 dao, sea salt was added daily to the pond (total 200 kg), but no reduction in mortality was observed. Subsequently, oxytetracycline (OTC) (200 mg/g active ingredient, Pharmatech, Thailand) was continuously administered to the fish via feed admixture (5g OTC per 1 kg of feed) for 7 days.

### 2.2. Gross necropsy and histopathology

Representative 10 fingerlings were collected at 6 dao for further examination approved by the Institutional Animal Care and Use Committee of Kasetsart University (Approval ID: ACKU63-FIS-009). After euthanasia with clove oil (150ppm/l), fresh skin mucus and gill samples were collected for microscopic examination. Bacteriological examination of brain, liver and kidney tissue samples was performed by streaking on tryptic soy agar and brain-heart infusion agar (BHIA) (Himedia, India) and incubation at 28°C for 48 hours. Samples from whole body fish (n=5) and detached gills (n=10) were fixed in 10% neutral buffered formalin and processed routinely for histology. Paraffin-embedded gills were sectioned at 5 μm, stained with haematoxylin and eosin and examined under a light microscope.

### 2.3. DNA extraction, 16S rRNA amplification and sequencing

Genomic DNA was isolated from infected gills using the Tissue Genomic DNA Mini kit (Geneaid, Taiwan) according to the manufacturer’s instructions. The presence of chlamydial DNA was first examined using primers 16SIGF (5’-CGGCGTGGATGAGGCAT-3’) and 16SIGR (5’-TCAGTCCCAGTGTTGGC-3’) described in the previous study [18]. All positive samples were subjected to further PCR with *Chlamydiales*-specific primers 16SIGF (5’-CGGCGTGGATGAGGCAT-3’) and 806R (5’-GGAC TACCAGGGTATCTAAT-3’) according to Relman [19]. PCR amplification reaction and cycling conditions for these assays were as previously described by Sood et al [20] and Draghi et al [21]. The expected amplicons of the first and second PCR methods were 300 bp and 766 bp, respectively. Amplified PCR products (766 bp) from each fish were isolated using NucleoSpin™ Gel PCR Clean-up Kit (Fisher Scientific, Sweden) according to the manufacturer’s protocol and then submitted for sequencing service (U2Bio, Thailand).

### 2.4. Phylogenetic analyses

A BLASTn query against available nucleotide sequences was deposited in the GenBank database (www.ncbi.nlm.nih.gov) to determine taxonomic identity. The closest known relatives and several sequences from related species of *Chlamydia*-like organisms were obtained from the NCBI database and used for phylogenetic analysis. The phylogenetic tree was constructed using the neighbour-joining method with 1,000 bootstraps after multiple alignments against the closely related *Chlamydia* bacteria using ClustalW in MEGA X version 10.2.4 [22]. To root the tree, sequences from a Betaproteobacterium, *Candidatus* Branchiomonas cysticola (Accession number JQ723599.1), were used as outgroup.

### 2.5. *In situ* hybridization

To confirm localization of CPT in the infected tissues, *in situ* hybridization (ISH) with a CLO-specific probe was performed on tissues from five representative diseased fish. A 766 bp probe was produced firstly by PCR using DNA extracted from infected fish as DNA template and the primers 16SIGF and 806R [19]. The product was labelled with digoxigenin (DIG) using a commercial PCR DIG labelling mixture (Roche Molecular Biochemicals, Germany) according to the manufacturer’s instructions. *In situ* hybridization was performed as previously described by Dinh-Hung et al [23] with some modifications. Briefly, unstained 4-μm sections on HistoGrip-coated slides (Fisher Scientific, Sweden) were deparaffinized 2 times for 5 minutes in xylene and then rehydrated through a graded series of ethanol and distilled water. After rapid treatment with cold acetic acid for 20 seconds and washing in distilled water, each section was covered with prehybridization buffer (4× SSC contain 50% [v/v] deionized formamide) for at least 10 minutes at 37°C. The probe was diluted in hybridization buffer (50% deionized formamide, 50% dextran sulphate, 50× Denhardt’s solution [Sigma-Aldrich, Germany], 20× SSC, 10 mg per ml salmon sperm DNA [Invitrogen, USA]), heated to 95°C for 10 minutes, and then cooled on ice. The specific probe was added to the slides, then covered with coverslips and incubated overnight at 42°C in a humidity chamber. For control slides, no probe was added to the hybridization solution. After hybridization, slides were washed in a series of graded sodium citrate solutions for 5 minutes in 2× SSC at room temperature (RT), 15 minutes in 2× SSC (37°C), 15 minutes in 1× SSC (37°C), 30 minutes in 0.5× SSC (37°C), and then equilibrated for 5 minutes in buffer I (1 M Tris-HCl, 1.5 M NaCl, pH 7.5). The tissue sections were then blocked with blocking solution buffer II (containing 0.1% Triton X-100 and 2% normal sheep serum) at room temperature for 30 minutes before being covered with anti-digoxigenin, Fab fragments (Roche Molecular Biochemicals, Germany, diluted 1:500 in buffer II) for 1 hour at 45°C. After washing twice for 10 minutes each with buffer I, sections were treated for 10 minutes in buffer III (100 mM Tris-HCl, 1.5 M NaCl, 50 mM MgCl_2_, pH 9.5). Signals were developed using the BCIP/NBT substrate, followed by counterstaining with nuclear fast red. The slides were then mounted, observed and photographed under a digital microscope.

## 3. Results

### 3.1. Initial diagnosis

Disease diagnosis carried out in the field on 6 dao showed no external lesions on the body surface of the fish **(Figure 1A)**. Wet-mount examination of the skin mucus and gills showed no external parasites, although the formation of several characteristic cysts in the gill filaments was prevalent **(Figure 1B)**. The infected fish exhibited lethargy, fish gasping at the water surface, and loss of appetite followed by mortality. This observation initially led us to a preliminary misdiagnosis as an “epitheliocystis” case. After treatment with OTC, there was a significant decrease in daily mortality of 150, 50 and 16 fish at 7, 8 and 9 dao, respectively. No mortality (0 dead fish) was observed after treatment with OTC for 4 days (10 dao).

**FIGURE 1.**
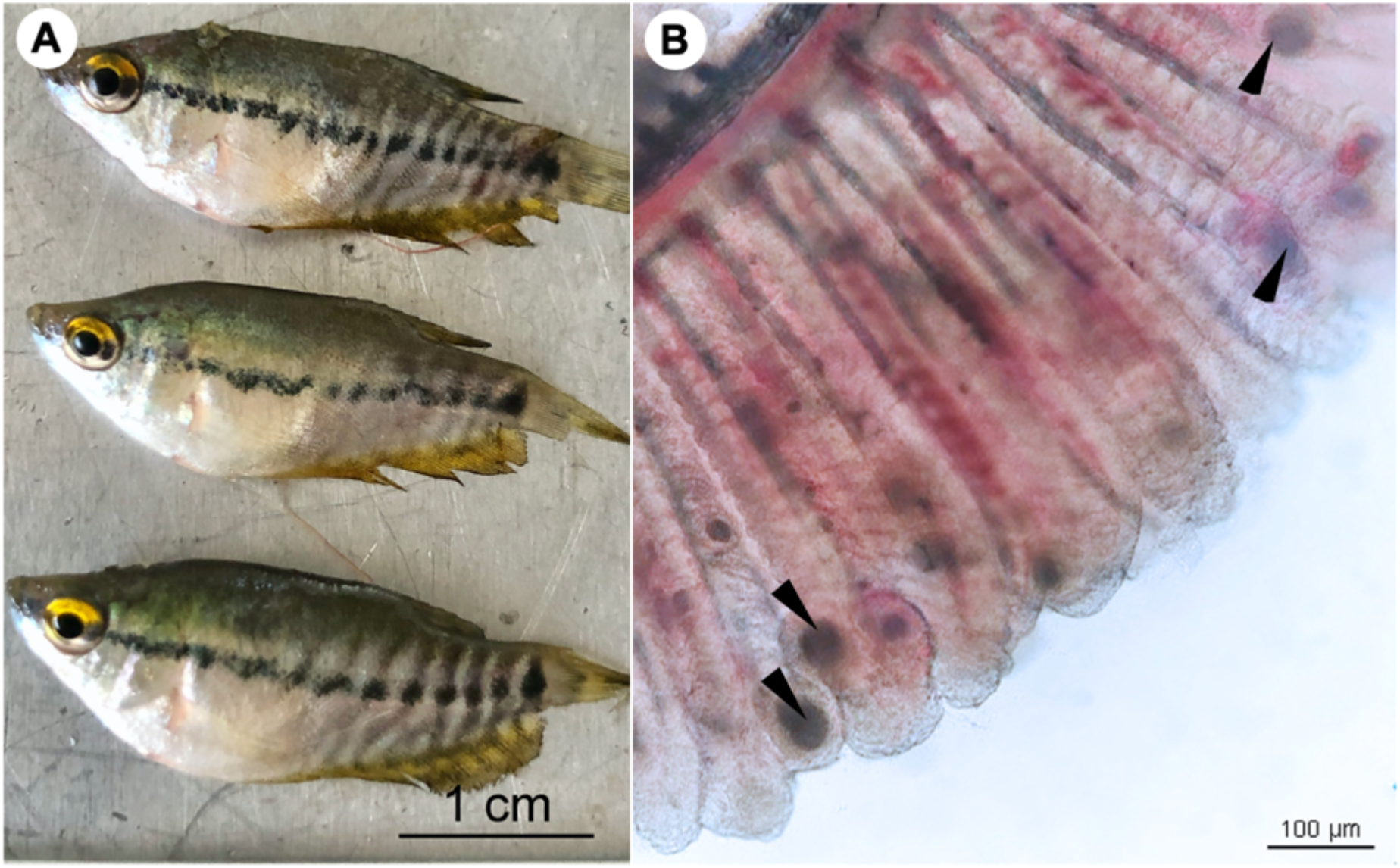
**(A)** Snakeskin gourami (*Trichopodus pectoralis*), no external lesions on the body surface of the fish. **(B)** Wet mount preparations of infected fish gill showing numerous morphological characteristics of “cysts” (arrowheads). Scale bars are shown in the pictures.

### 3.2. Histopathology and *in situ* hybridization

Histological examination revealed colonization by intracellular bacteria in several organs, including the gill filaments **(Figure 2A.1)**, submucosa of the intestine **(Figure 2B.1)**, and caudal fin tissue **(Figure 2C.1)**. Dense, roundish to oval intracellular bacteria with a great affinity for connective tissue rather than epithelial cells were observed **(Figure 2A.2, B.2, and C.2)**. Apparently, the bacteria infected the primary gill filaments rather than the secondary gill filaments. In particular, the cartilaginous junctions between primary and secondary gill filaments were apparently more susceptible to infection than others, resulting in the separation of the two components **(Figure 2A.2)**. Similar changes were also observed at the cartilaginous junction of the caudal fin (data not shown). With respect to ISH, the DIG-labeled probe was specifically bound to intracellular bacterial foci **(Figure 2A.3, B.3, and C.3)**, whereas no binding signal was observed in tissue sections incubated with no probe **(Figure 2A.4, B.4, and C.4)**. Furthermore, no culturable bacteria were isolated from the internal organs on nutrient agar plates as well as on brain-heart infusion agar plates even after 48 hours of incubation.

**FIGURE 2.**
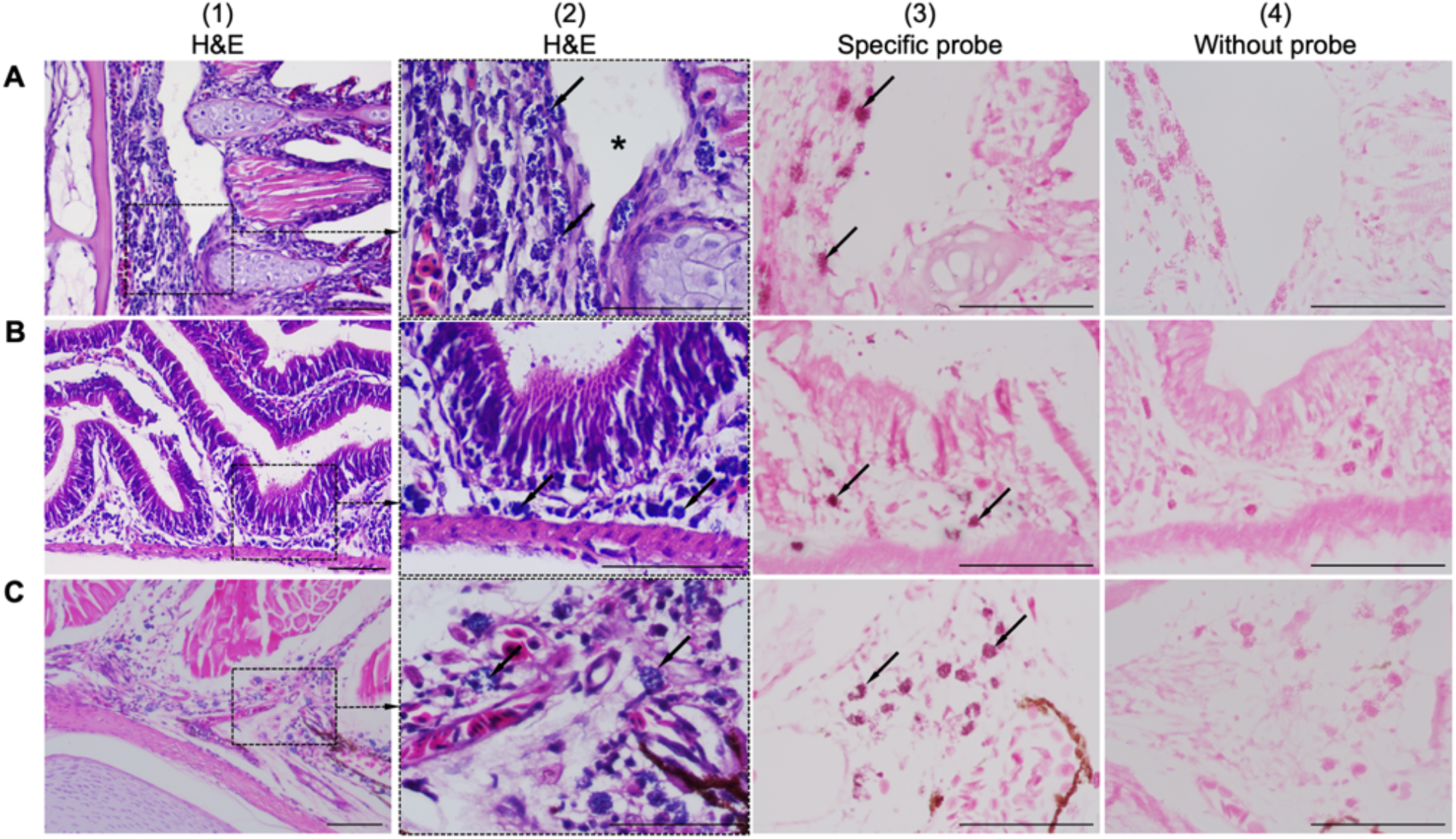
Comparison of consecutive gill sections of infected fish stained with H&E **(A.1 - C.1; A.2-C.2)**, ISH with a specific probe **(A.3-C.3)** and ISH without probe **(A.4-C.4)** as control. Infected fish showed the presence of a novel intracellular bacterium in the gill **(A.1)**, intestinal submucosa **(B.1)** and caudal/tail fin **(C.1)**. Higher magnification showed that the connective tissue was more susceptible to infection than others due to colonization by dense, roundish to oval, blue-stained intracellular bacteria (arrows in Figure **A.2, B.2**, and **C.2**). The bacteria were observed near the cartilaginous junctions of the primary and secondary gill filaments, resulting in disruption of the tissue junction (asterisk in Figure **A.2**). ISH positive reactivity of intracellular bacterial foci is indicated by distinct dark signals (arrows in Figure **A.3-C.3**). Scale bar = 50 µm.

Interestingly, histological analysis showed that the “cysts” found in the infected fish were not epitheliocystis, as tentatively diagnosed and later identified as plasmodia of a myxosporean. The presence of numerous plasmodia was observed on the gill filaments **(Figure 3A)**. Myxospores and plasmodia demonstrated asynchronous development with young round plasmodia encased in a wall of epithelial cells of the gill filaments **(Figure 3B)**. As the plasmodium grew, the envelope ruptured and released myxospores into the adjacent tissue **(Figure 3C)**. The myxospores exhibited typical features of the genus *Henneguya*: two equal polar caps, sporoplasm at the posterior pole of the spore, and two long, superimposed caudal processes **(Figure 3D)**. Histological analysis of the infected gills of *Henneguya* showed that the plasmodia caused severe distortion of the lamellar structure and obstruction of the gills by compression of the cysts **(Figure 4A, B, and C)**. The plasmodia occupied the part extent of the gill lamellae and produced marked dilatation and discrete epithelial hyperplasia **(Figure 4D and E)**. The extensive dilatation of the infected lamellae caused displacement and deformation of the adjacent lamellae. As the plasmodia grew, they compressed the adjacent tissue and caused tissue necrosis in the infected area **(Figure 4E and F)**.

**FIGURE 3.**
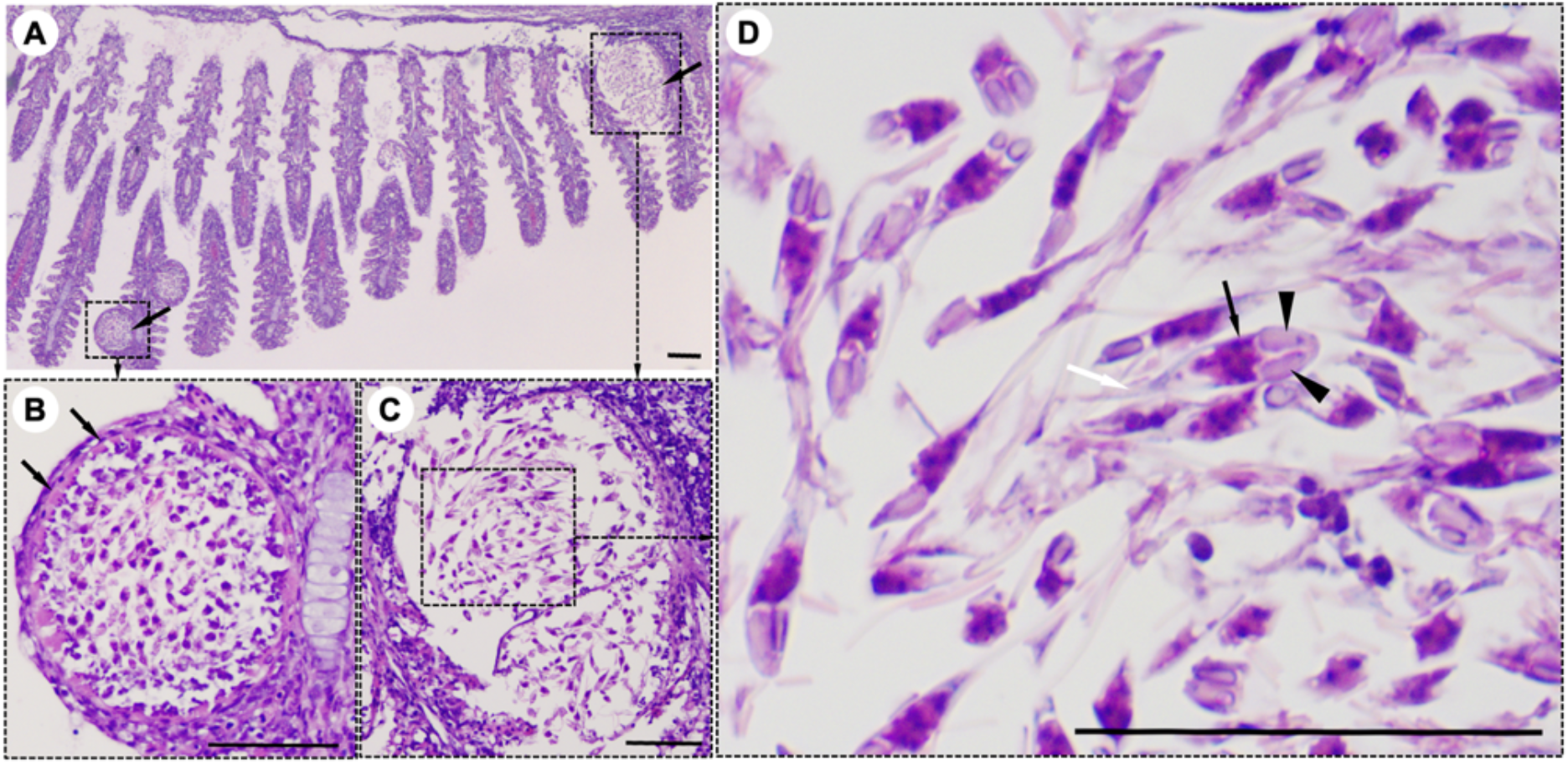
Histological lesions in the gills of the snakeskin gourami (*Trichopodus pectoralis*) infected with *Henneguya* sp. **(A)** Plasmodia (arrows) showing different developmental stages. **(B)** Young plasmodium was roundish and encased in a wall of epithelial cells (arrows). Immature myxospores were located in the periphery of the plasmodia and mature myxospores in the centre. **(C)** A grown plasmodium ruptured the envelope. **(D)** The myxospores have typical features of a *Henneguya* sp. including two equal polar caps (arrowheads), sporoplasm at the posterior pole of the spore (black arrow), and two long, superimposed caudal processes (white arrow). Slides were stained with hematoxylin and eosin (H&E), scale bar = 50 µm.

**FIGURE 4.**
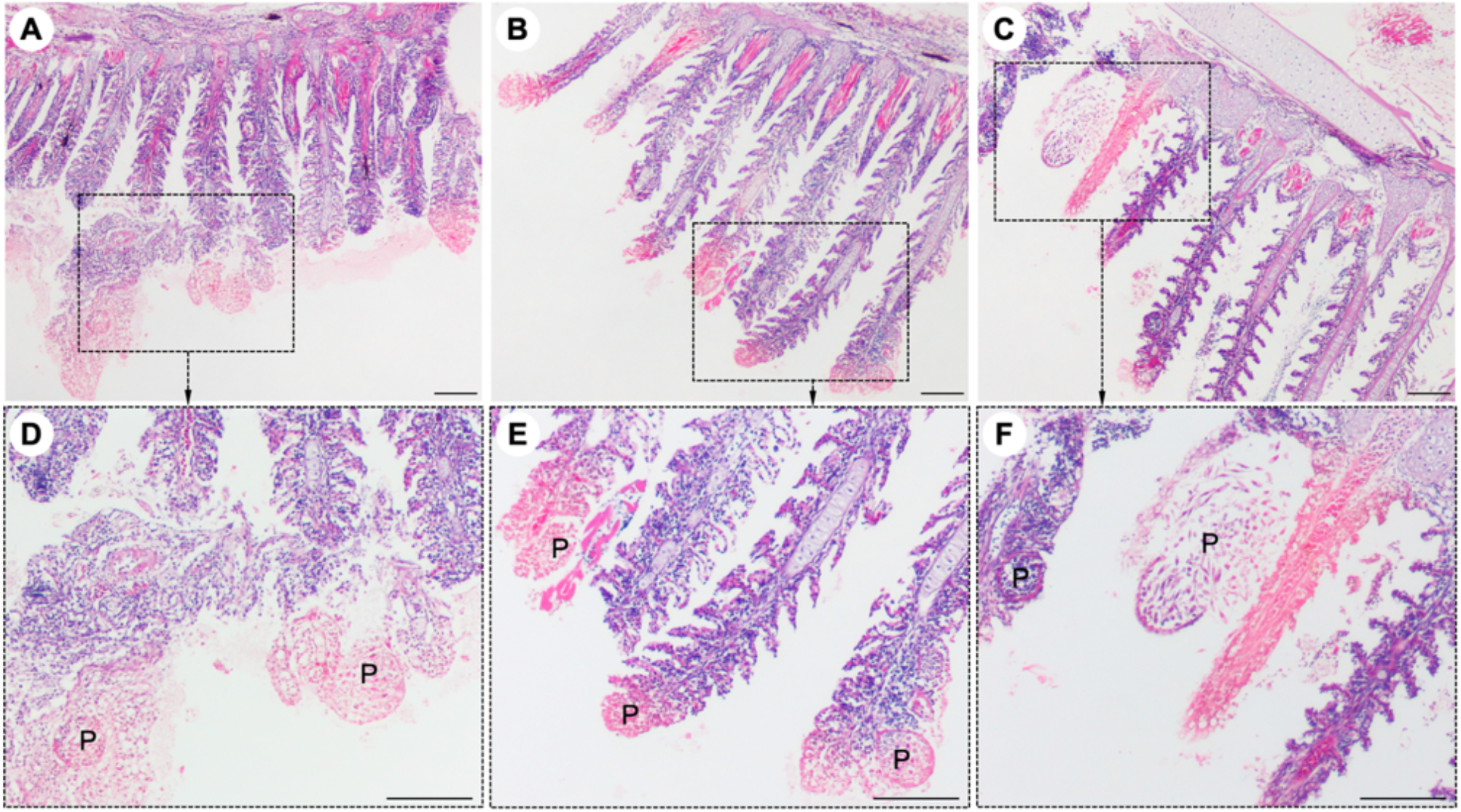
Histological lesions in the gills of the snakeskin gourami (*Trichopodus pectoralis*) infected with *Henneguya* sp. The plasmodia (P) caused severe distortion of the lamellar structure and obstruction of the gills by compression of the cysts **(A, B, C)**. Higher magnification indicated plasmodia occupied part of the gill lamellae and caused marked dilation and discrete epithelial hyperplasia **(D, E)**. The plasmodia grew and compressed the adjacent tissue and caused tissue necrosis in the infected area **(E, F)**. Slides were stained with hematoxylin and eosin (H&E), scale bar = 100 µm.

### 3.3. Molecular analyses

All collected samples (n=4) were positive after repeated PCR amplification of the 16S rRNA gene of *Chlamydiales* **(Figure S1)**. Multiple sequence alignment of the partial 16S rRNA gene sequence of the *T. pectoralis* chlamydial pathogen from 4 representative infected fish, showed that the sequences were identical, and the consensus sequence (766bp) was deposited in GenBank under accession number MW832782. BLAST-n search of consensus sequence in the NCBI database revealed closest sequence similarity (93.5%) with the partial 16S rRNA gene of a non-cultured bacterium (Accession number LN612734.1) obtained from a case of gill disease in Mediterranean Sea bream, followed by *Candidatus* Piscichlamydia sp. (92.3%) (Accession number KY380090.1), which is associated with epitheliocystis infections in cyprinids. These bacteria formed a clade as an unclassified family belonging to the genus *Candidatus* Piscichlamydia within the order *Chlamydiales*. The accession numbers and taxonomic identities, as well as the origin of the organisms included in this phylogenetic analysis, are shown in **Figure 5**. This phylogenetic analysis confirmed that the bacterial pathogen in the case of the snakeskin gourami separated in a unique branch, representing a novel species, proposed name “*Candidatus* Piscichlamydia trichopodus”, a new member of the order *Chlamydiales*.

**FIGURE 5.**
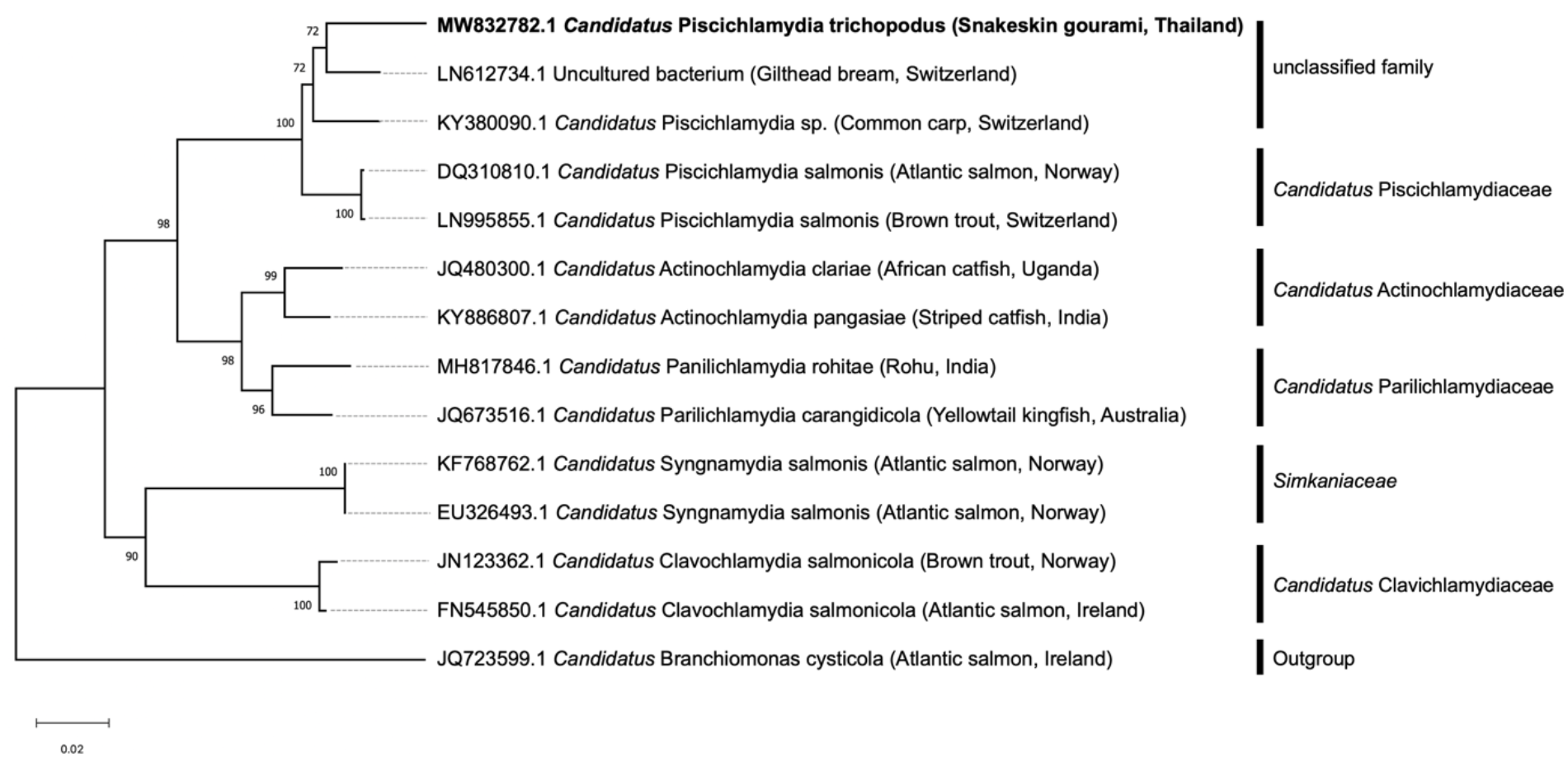
The phylogenetic tree was constructed based on the partial 16S rRNA sequence (766 bp) of the snakeskin gourami (*Trichopodus pectoralis*) from this study (MW832782) and closely related species. The accession numbers and taxonomic identities, as well as the host origin of the organisms included in this phylogenetic analysis, are shown. *Candidatus* Branchiomonas cysticola was selected as the outgroup. The tree was constructed using the neighbour-joining method. The scale bar represents 0.02 - nucleotide substitution per site, while the number at the node of the tree indicates the bootstrap value in percent.

## 4. Discussion

Members of the order *Chlamydiales* are a diverse group of Gram-negative, obligate intracellular bacteria that are distributed worldwide and cause a wide variety of diseases in humans, livestock, domestic animals, wildlife, and exotic species [16, 24, 25]. *Chlamydia-*like organisms (CLOs), commonly found in aquatic environments, have been identified as causing disease in at least 90 fish species in both freshwater and marine environments [14-16]. A common feature of these bacteria is the infection of epithelial cells of fish, causing typical lesions in the form of epitheliocystis [13-16, 26, 27]. Epitheliocystis as a result of CLOs infection have been described in several farmed fishes, including common carp *Cyprinus carpio* [28], red seabream *Pagrus major* [13], African catfish *Clarias gariepinus* [29], yellowtail kingfish *Seriola lalandi* [30], striped trumpeter *Latris lineata* [31] barramundi *Lates calcarifer* [32], striped catfish *Pangasius hypophthalmus* [33], and rohu *Labeo rohita* [20]. In contrast to previous studies, our result showed that the histopathological changes associated with CPT infection revealed massive intracellular colonization but not obvious as epitheliocystis. Interestingly, the microbial pathogen in this study likely shares similarities with the CLO pathogen that causes systemic microbial disease in the Dungeness crab, *Cancer magister* [34]. Both pathogens show systemic infection with numerous colonies of the organism having strong affinity for connective tissue and connective tissue cells, while rarely infecting epithelial cells. Several previous studies reported that CLOs can infect other cell types, including mucous cells in common carp *Cyprinus carpio L*. [35], pillar cells in tiger puffer *Takifugu rubripes* [36], macrophages in brown bullhead *Ictalurus nebulosus* [37] and chloride cells in Atlantic salmon *Salmo salar* [38]. These studies have also indicated that epitheliocystis due to CLO is typical but not always observed. Moreover, the localization of CPT near the damaged cartilaginous tissue suggests that these bacteria may require cartilage for their metabolism. This histopathological feature could be considered in the presumptive diagnosis of this new pathogen. It is increasingly recognized that these pathogens are actually diverse in their morphology, e.g., the morphology of CLO organisms in the epithelial cysts and their staining characteristics, as well as the morphology of their capsules and their location in fish tissue [39-41]. This is the first detection of CLO in snakeskin gourami and may represent another *Chlamydia* that is not associated with epitheliocystis. Since CLO pathogens cannot be distinguished by morphology or conventional culture methods, molecular methods are widely used to detect and characterize the causative pathogen. Apart from histopathological lesion of the bacterial foci, CLO infection was confirmed by ISH and specific PCR assay. Although the short 16S rRNA signature sequence detected in infected tissue is not ideal for detailed phylogenetic analysis, it is unique. Our sequence showed only a distant similarity of 93.5% and 92.3%, respectively, with a query coverage of 98% to the uncultured bacterium from gilthead sea and *Candidatus* Piscichlamydia sp. associated with epitheliocystis infections in cyprinids [27]. There bacteria formed a clade as an unclassified family belonging to the genus *Candidatus* Piscichlamydia within the order *Chlamydiales* and also shared >90% sequence similarity with other members of the family *Candidatus* Piscichlamydiaceae, suggesting that these pathogens belong to the same family [18] or are closely related. Since the causative agent shared < 95% similarity with other previously reported *Chlamydia-*like 16S rRNA sequences, the sequenced bacteria are new to the order *Chlamydiales* [18]. These results highlight the wide genetic diversity within this bacterial group and are consistent with previous findings [26] that each new fish host indicates the existence of a phylogenetically distinct and novel *Chlamydia* infection.

In the present case, the diagnosis was challenging due to mixed infection of CPT along with a *Henneguya* sp parasite in the affected gill tissues. Myxospore taxonomy is based primarily on morphology and spore structure, therefore, the myxosporids found in this study can be morphologically classified into the genus *Henneguya* based on certain morphological criteria. *Henneguya* sp. has the characteristic morphological feature of two equal polar caps, sporoplasm at the posterior pole and 2 independent caudal processes, which distinguishes it from the other genus of the family *Myxobolidae* [7, 42, 43]. The species of the genus *Henneguya* interact with the gill structures of fish in different manners, resulting in varying degrees of disease [9, 10, 44]. The clinical signs noted by the farmer in the snakeskin gourami, particularly lethargy, gasping for oxygen at the water surface, and loss of appetite, are comparable with those previously documented in fish affected by *Henneguya* sp. [45-47]. The most common histological lesions found in this case were distortion of the lamellar structure and obstructions of the gills due to compression of the cysts, which most likely caused respiratory distress and contributed to the observed mortality. Histopathological features similar to those described in this study have been observed in previous studies [48-50], in which parasitism by *Henneguya* sp. resulted in marked dilatation and discrete epithelial hyperplasia, and continued growth of the parasitic cyst resulted in tissue necrosis in the surrounding infected area. The observation of areas of cystic lesions associated with tissue necrosis caused by the compression of cysts in the epithelial cells of the gill lamellae in this study has also been reported in previous studies [51]. The growth of *Henneguya* plasmodia results in a displacement and distortion of the lamellar structures, possibly adversely affecting gas exchange and associated with mortality in the fish population

Unfortunately, due to the occurrence of co-infections and the unavailability of culture methods for fish CLOs, it was not possible to determine which pathogens were directly responsible for the associated mortality. Interestingly, the disease is sensitive to tetracyclines, since the antibiotic treatment with oxytetracycline in the field was effective, hence we assumed that the disease might have been caused primarily by the CLO. As previously reported by Goodwin et al. [52], OTC significantly reduced mortality due to chlamydial infection and supported this treatment regimen for future outbreaks. The cause and pathogenesis of this dual infection remain speculative and await the development of a culture technique and isolation of the *Chlamydia*-like organism before further *in vitro* studies. It is unclear whether *Henneguya* sp. and CPT has the potential of causing a significant impact on snakeskin gourami aquaculture. Given the commercial importance of the snakeskin gourami *Trichopodus pectoralis* and its great aquaculture potential, the results of this study highlight the need of follow up investigations on ultrastructural morphology, host range, prevalence, risk factors for disease, and mode of transmission. This could lead to a better understanding of the pathogen’s biology and disease epidemiology for development of effective control measures.

In conclusion, the presence of pathogenic potential of mixed infection of a novel intracellular CLO (*Candidatus* Piscichlamydia trichopodus) and a gill parasite *Henneguya* sp. in snakeskin gourami in Thailand is reported for the first time. This study expands the knowledge of the pathology of snakeskin gourami, an important fish species in Asian aquaculture, and contributes to initial understanding of diseases in this fish species.

## ACKNOWLEDGEMENTS

The work was carried out with support from the Center of Excellence in Aquatic Animal Health Management, Faculty of Fisheries, Kasetsart University.

## Supplementary data

**FIGURE S1.**
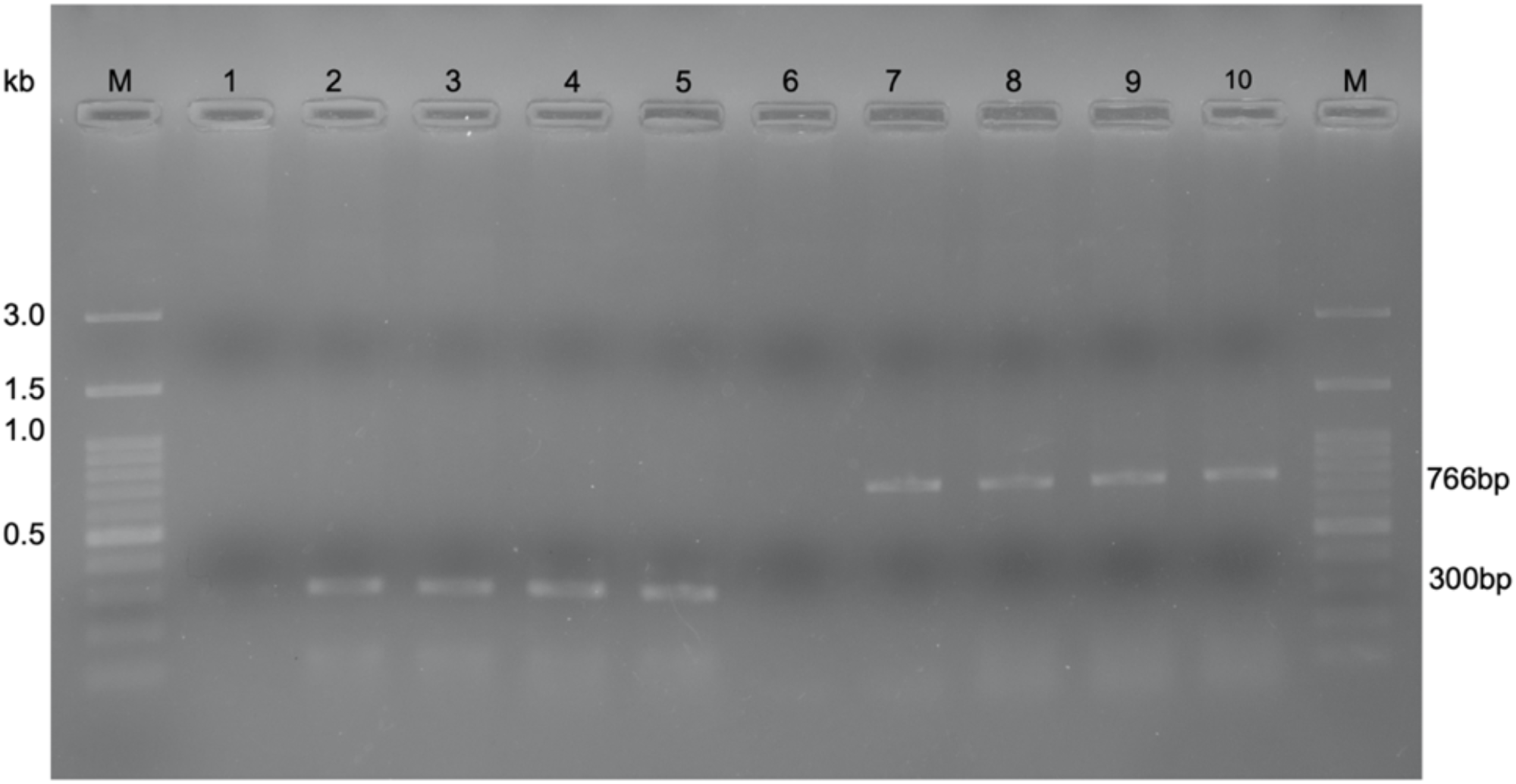
Confirmation of PCR test results under agarose gel electrophoresis of samples from four representative fish. Lanes 1 and 6 were amplified without DNA template using primer set 1 and 2, respectively, as negative controls. Lanes 2, 3, 4 and 5 were amplified with the DNA template from primer set 1 (300bp). Lanes 7, 8, 9 and 10 were amplified with the DNA template from primer set 2 (766bp). Lane M was a DNA marker (Himedia, India).

